# The kinetics of ribonucleoprotein granules

**DOI:** 10.64898/2026.02.04.703460

**Authors:** Yuchen Li, Xiangting Li, Mengmeng Xu, Zhi Qi

**Affiliations:** Center for Quantitative Biology, Academy for Advanced Interdisciplinary Studies, Peking University, Beijing 100871, China; Peking-Tsinghua Center for Life Sciences, Academy for Advanced Interdisciplinary Studies, Peking University, Beijing 100871, China; Department of Computational Medicine, University of California, Los Angeles, CA 90095, USA; Tsinghua-Peking Center for Life Sciences, Tsinghua University, Beijing 100084, China; School of Physics, Peking University, Beijing 100871, China

## Abstract

Ribonucleoprotein (RNP) granules are biomolecular condensates composed of diverse RNAs and RNA-binding proteins that play essential roles in regulating multiple aspects of RNA biology. As complex biological systems, the condensate functions of RNP granules emerge from the higher-order kinetic assembly of individual molecules. Here, we develop an in vitro single-molecule approach—termed SMART (group single-molecule assay for ribonucleoprotein granules)—that enables direct and quantitative measurements of RNP granule assembly and disassembly kinetics across nanoscale to mesoscale regimes. Using SMART, we reveal that RNP granule assembly is strongly pathway dependent. In parallel, we develop a minimal two-state mathematical model that faithfully reproduces these kinetic behaviors. This model enables quantification of multivalent interaction timescales, reconstruction of the underlying free-energy landscape governing RNP granule dynamics, and demonstrates that this assembly process is intrinsically non-equilibrium. Understanding the condensate functions of RNP granules suggests new strategies for their rational manipulation in health and disease.

## Introduction

Ribonucleoprotein (RNP) granules (*1*) are biomolecular condensates composed of diverse RNAs and RNA-binding proteins (RBPs), and they play crucial roles in regulating various aspects of RNA biology. These granules are multicomponent assemblies formed through multivalent and weak interactions, including RNA-RBP binding as well as homotypic and heterotypic interactions among RBPs and RNAs. Dysregulation of RNP granules has been implicated in numerous human diseases, underscoring their importance in cellular physiology.

RNP granules constitute a class of mesoscale complex biological systems whose structures and functions emerge from nanometer-scale biomolecular interactions to mesoscale organization (*2, 3*). Interestingly, the length of RNA substrates within RNP granules has been shown to influence their assembly dynamics; here, we focus on RNP granules formed on long RNAs, such as cytoplasmic granules containing long messenger RNAs. A central unresolved question is how the condensate functions of RNP granules arise from the higher-order assembly of individual molecules. Although extensive studies have established that both homotypic and heterotypic interactions collectively shape granule composition (*4–7*), experimental tools of the kinetics governing this cross-scale assembly process remains challenging. This challenge stems from the diversity of interacting molecular species and the complexity of their multivalent interactions.

To address these gaps, we established a novel single-molecule approach termed the group <u>s</u>ingle-<u>m</u>olecule <u>a</u>ssay for <u>r</u>ibonucleoprotein granules (SMART). Built upon the high-throughput DNA Curtains platform (*8–10*), SMART enables direct, quantitative measurements of RNP granule assembly and disassembly kinetics across nanoscale and mesoscale regimes. In parallel, we developed a two-state mathematical model that accurately simulates these kinetic behaviors and yields a free-energy landscape describing RNP granule dynamics. By extending classical physical chemistry frameworks used to describe the kinetics of elementary chemical reactions (*11*), our approach provides a quantitative framework for dissecting the whole kinetic principles governing biomolecular condensates, which contains large amount of elementary chemical reactions above in physical chemistry.

We applied SMART in combination with the mathematical model to investigate RNP granules composed of either a single RBP (single-component RNP granule) or two distinct RBPs (two-component RNP granule). Our results demonstrate that the assembly kinetics of single-component RNP granules are intrinsically non-equilibrium, whereas the assembly of two-component granules is strongly pathway dependent. Collectively, SMART and the accompanying theoretical framework provide a quantitative platform for elucidating multiple biophysical condensate functions of RNP granules. More broadly, these insights may ultimately enable the precise manipulation of RNP granule dynamics, opening new avenues for therapeutic intervention in human disease.

## Results

### Developing group <u>s</u>ingle-<u>m</u>olecule <u>a</u>ssay for <u>r</u>ibonucleopro<u>t</u>ein granules (SMART)

To monitor the cross-scale kinetics of RNP granules, we first inserted a single T7 promoter into Lambda DNA and introduced these DNA substrates into a microfluidic flow chamber of the DNA Curtains platform (Supplementary Fig. 1) (*8–10*). Individual DNA molecules were tethered to supported lipid bilayers via specific streptavidin-biotin interactions. T7 RNA polymerase (T7 RNAP) and a mixture of nucleoside triphosphates (NTPs) and Fluor647X-labeled UTP, were then introduced to generate nascent RNA transcripts from identical DNA templates. These RNAs were visualized as localized Fluor647X-enriched regions downstream of the T7 promoter and extending toward the DNA terminus (Fig. 1a(i)-(ii), b(i) and (vi)). Quantitative analysis of RNA transcript length (nucleotides, nts), inferred from Fluor647X fluorescence intensity, yielded an average of (4.1 ± 3.9) × 10^5^ nts (mean ± s.d., N = 1,455, Supplementary Fig. 2a(i)). Interestingly, the RNA transcripts remained stably associated with the Lambda DNA templates (*12*). This persistent association was further confirmed using a two-color in vitro transcription assay (Supplementary Fig. 3).

**Fig. 1.**
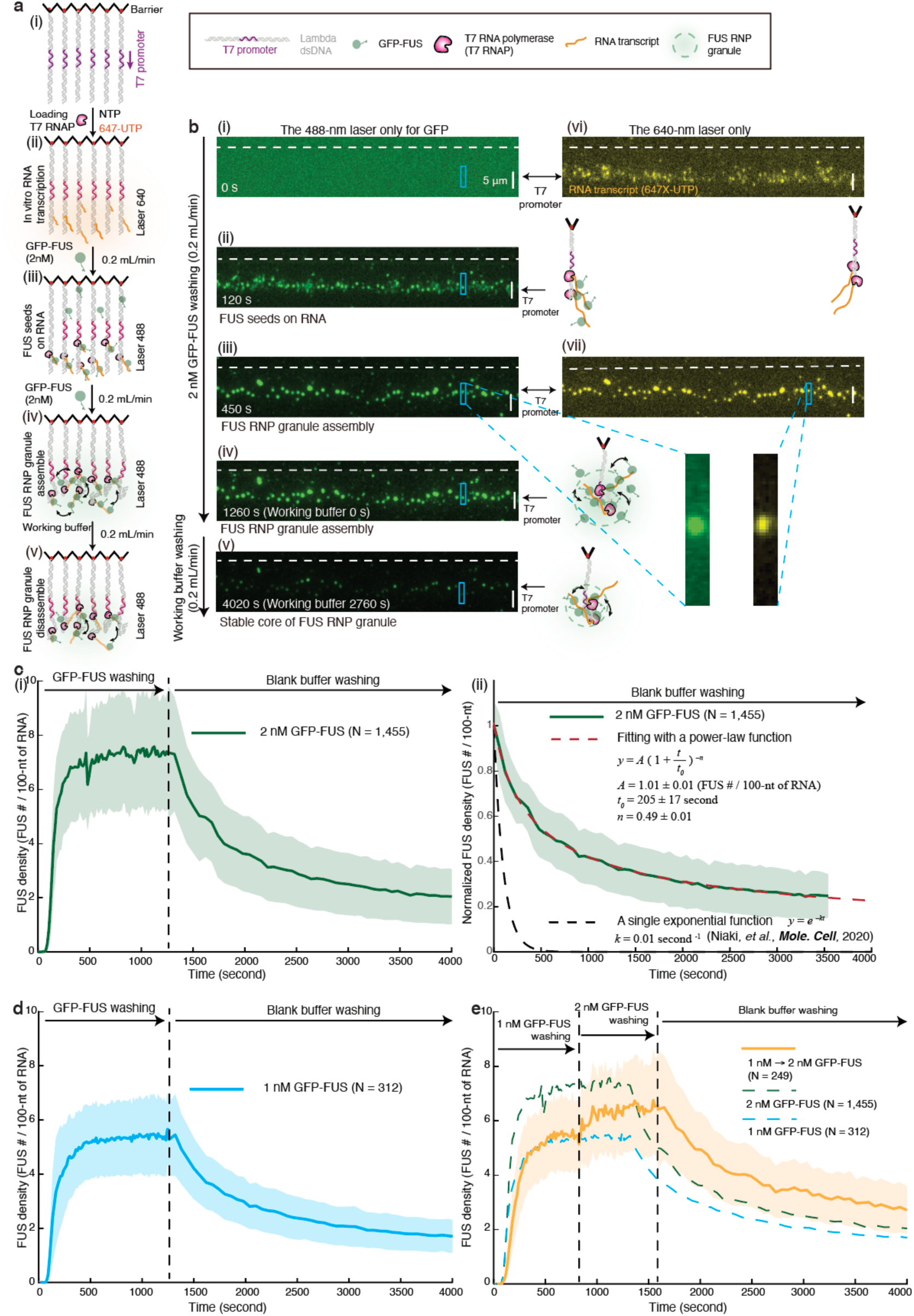
Quantitative measurements of FUS RNP granule assembly and disassembly kinetics using SMART. (**a**) Schematic illustrating the SMART strategy for quantifying FUS RNP granule kinetics. (**b**) Wide-field total internal reflection fluorescence (TIRF) microscopy images acquired during SMART experiments with 2 nM FUS during the assembly phase (i)-(iv), followed by washing with blank buffer during the disassembly phase (v). Panels (vi)-(vii) show the corresponding RNA fluorescence signals. (**c**) (i) SMART measurement of granule assembly dynamics during exposure to 2 nM FUS for 1,260 seconds, followed by washing with blank working buffer. (ii) Normalized dissociation curve derived from (i) upon blank-buffer washing. The black dashed line indicates a single-exponential decay obtained from single-molecule measurements of individual FUS dissociation, whereas the red dashed line shows a power-law fit to the SMART data. N = 1,455, which was the total puncta number of FUS RNP granules examined over nine times SMART experiments (n = 9). Data are presented as mean ± s.d. (**d**) SMART measurement of granule assembly dynamics during exposure to 1 nM FUS for 1,260 seconds, followed by washing with blank working buffer. N = 312, which was the total puncta number of FUS RNP granules examined over two times SMART experiments. Data are presented as mean ± s.d. (**e**) Assembly kinetics measured under sequential protein exposure (Orange line), in which the chamber was first equilibrated with 1 nM FUS for 825 seconds and subsequently exposed to 2 nM FUS for an additional 765 seconds. N = 249, which was the total puncta number of FUS RNP granules examined over two times SMART experiments. Data are presented as mean ± s.d. The green and blue dashed lines are SMART measurements of Fig. 1c(i) and d. The working buffer for SMART is 40 mM Tris-HCl (pH 7.5), 75 mM KCl, 0.5 mg/mL BSA, 2 mM MgCl_2_, and 1 mM DTT, and 8 unit/mL RNasin® Plus RNase Inhibitor.

To verify successful transcription of RNA substrates within the flow cell, we cloned DNA sequences encoding either 8× Pepper or 6× MS2 stem-loop arrays downstream of the T7 promoter on Lambda DNA. Upon introduction of the fluorescent dye HBC620 (*13*) or purified mCherry-labeled MS2 coat protein (MCP) (*14*), robust fluorescent signals corresponding to HBC-Pepper or MCP-MS2 interactions were detected within the flow cell (Supplementary Fig. 4). These observations confirm efficient in vitro transcription in the flow cell and further indicate that the local RNA secondary structures required for RBPs binding are properly formed and stably maintained.

Once RNA substrates were generated within the flow chamber, 2 nM GFP-tagged FUS (GFP-FUS; Supplementary Fig. 5) was introduced at a flow rate of 0.2 mL/min. We observed the rapid emergence and growth of green fluorescent puncta downstream of the T7 promoter (Fig. 1a(iii)-(iv), b(ii)-(iv)), concomitant with progressive compaction of the RNA substrates (Fig. 1b(vi) and (vii)). For quantitative analysis, fluorescence signals from nascent RNA transcripts were combined with calibrated GFP intensities to estimate protein density (Supplementary Fig. 6-7 and **Supplementary Methods**). This analysis enabled calculation of the number of FUS molecules per 100-nt of RNA, defined here as the FUS density within RNA-FUS clusters (Fig. 1c(i)). FUS assembly exhibited a clear saturation phase, indicative of the formation of stable RNA-FUS clusters. At saturation (1,260 seconds), the FUS density reached 7.3 ± 2.0 (mean ± s.d., N = 1,455, Supplementary Fig. 2b), corresponding to an average of (3.0 ± 3.1) × 10^4^ FUS molecules per cluster (mean ± s.d., N = 1,455; Supplementary Fig. 2a(ii)). The mean diameter of RNA-FUS clusters was 1.1 ± 0.6 μm (mean ± s.d., N = 1,455; Supplementary Fig. 2c-d). Following saturation, the chamber was continuously washed with blank working buffer. After 2,760 seconds of blank buffer washing, the FUS density decreased to 2.0 ± 1.0 (mean ± s.d., N = 1,455, Supplementary Fig. 2b), corresponding to an average of (1.0 ± 1.3) × 10^4^ FUS molecules per cluster (mean ± s.d., N = 1,455, Supplementary Fig. 2a(iii)), reflecting progressive disassembly of the RNA-FUS clusters (Fig. 1a(v) and b(v)).

Together, these experiments demonstrate that the RNA-FUS clusters—hereafter referred to as FUS RNP granules (Single-component RNP granules)—exhibit well-defined assembly and disassembly kinetics captured by the time traces in Fig. 1c(i). The assembly curve bridges nanometer-scale molecular interactions (RNA-protein, protein-protein, and RNA-RNA) with micrometer-scale RNP condensates, revealing the cross-scale nature of granule formation for the first time. This framework offers a direct and quantitative means to measure RNP granule kinetics, a method we term the group <u>s</u>ingle-<u>m</u>olecule <u>a</u>ssay for <u>r</u>ibonucleoprotein granules (SMART).

To validate this approach, we performed two control experiments. First, we characterized the photobleaching behavior of Fluor647X-labeled UTP and GFP under 640-nm and 488-nm illumination (Supplementary Fig. 8 and **Supplementary Methods**), confirming that RNA fluorescence within FUS RNP granules (Fig. 1b) was not substantially affected by photobleaching. Second, following transcription by T7 RNAP (Fig. 1b(vi)), nascent RNA transcripts remained stably inside FUS RNP granule tethered to their DNA templates for more than 5,000 seconds (Supplementary Fig. 9a), exceeding the full experimental timescale in Fig. 1b.

### FUS RNP granule possesses a stable core architecture

The disassembly kinetics in Fig. 1c(i) reveal a pronounced slowdown in FUS dissociation after approximately 2,760-s blank buffer washing, indicating the emergence of a stable core architecture within the FUS RNP granule. After approximately 2,760-s blank buffer washing, ∼67% of the FUS molecules present at saturation (1,260 seconds) had dissociated, whereas only ∼4% of the RNA substrates were released over the same interval (Supplementary Fig. 9b). By contrast, in the absence of FUS, ∼42% of RNA substrates dissociated from Lambda DNA under identical conditions (Supplementary Fig. 9a). Together, these observations support three key conclusions. First, the stable core is characterized by a low protein density (∼2 FUS molecules per 100 nt of RNA, Supplementary Fig. 2b), consistent with in vivo estimates for stress granules (∼0.6-1.5 proteins per 100-nt of RNA) (*1, 15, 16*); Second, despite substantial protein loss, the core structure retains more than 96% of the RNA substrates, suggesting that the RNA adopts specific secondary or higher-order architectures that stabilize the assembly (*17*). Consistent with this interpretation, treatment with RNase A, which degrades both single- and double-stranded RNA, resulted in digestion of ∼96% of RNA substrates within FUS RNP granules within 370 seconds. In contrast, RNase I_f_, which preferentially cleaves single-stranded RNA, degraded only ∼7% of RNA substrates within FUS RNP granules even after 3,800 seconds (Supplementary Fig. 10); Third, the assembly of FUS RNP granules is an irreversible process, and the dissociation of proteins does not lead to the spontaneous disassembly of the RNA structure.

### The assembly of FUS RNP granule is concentration and pathway dependent

We next reduced the protein concentration from 2 nM to 1 nM and repeated the experiment shown in Fig. 1b, with the results plotted in Fig. 1d. Under this lower concentration, both the assembly (5.3 ± 1.4 (mean ± s.d., N = 312 at 1,260 seconds)) and disassembly (1.7 ± 0.6 (mean ± s.d., N = 312 after 2,760 seconds of blank buffer washing)) reached a reduced saturation level compared to the 2 nM condition (Fig. 1d and Supplementary Fig. 11a), indicating that both assembly and disassembly kinetics of FUS RNP granules are regulated by protein concentration.

When the system was first equilibrated at 1 nM and subsequently exposed to 2 nM protein, the saturation level increased from 5.5 ± 1.3 (mean ± s.d., N = 249 at 825 seconds) to 6.6 ± 1.8 (mean ± s.d., N = 249 at 1,590 seconds) (Fig. 1e and Supplementary Fig. 11b-c). This finding suggests that not all RNA regions are occupied within the FUS RNP granule in 1 nM condition, strongly suggesting the existence of “accessibility” inside RNP granules. Interestingly, the saturation level of this new experiment (∼6.6, 1 nM → 2 nM GFP-FUS) is lower than ∼7.3 of pure 2 nM GFP-FUS washing experiment in Fig. 1c(i). This result suggests that the assembly kinetics of FUS RNP granule is pathway dependent. Furthermore, the saturation level of disassembly process in this exchange experiment (Fig. 1e) after 2,760-second blank buffer washing was approximately 2.6 ± 0.8 (mean ± s.d., N = 249, Supplementary Fig. 11d), which is higher to the value observed under continuous 2 nM FUS washing in Fig. 1c(i), suggesting that the stable core structure of the FUS RNP granule is also pathway dependent.

### A mathematical model reveals multiple interactions inside FUS RNP granules

We observed that a subset of FUS molecules within FUS RNP granules remained bound to RNA for over 3,600 seconds, and the disassembly kinetics deviated from a single-exponential decay. Instead, the decay followed a power-law scaling (Fig. 1c(ii)), consistent with a broad distribution of dissociation rates (*18, 19*). These kinetics imply that RNP granules sample multiple interaction states—encompassing RNA-protein, protein-protein, and RNA-RNA contacts—thereby motivating the development of a mathematical model (Fig. 2a and **Supplementary Methods**) to interpret the kinetic features revealed by SMART.

**Fig. 2.**
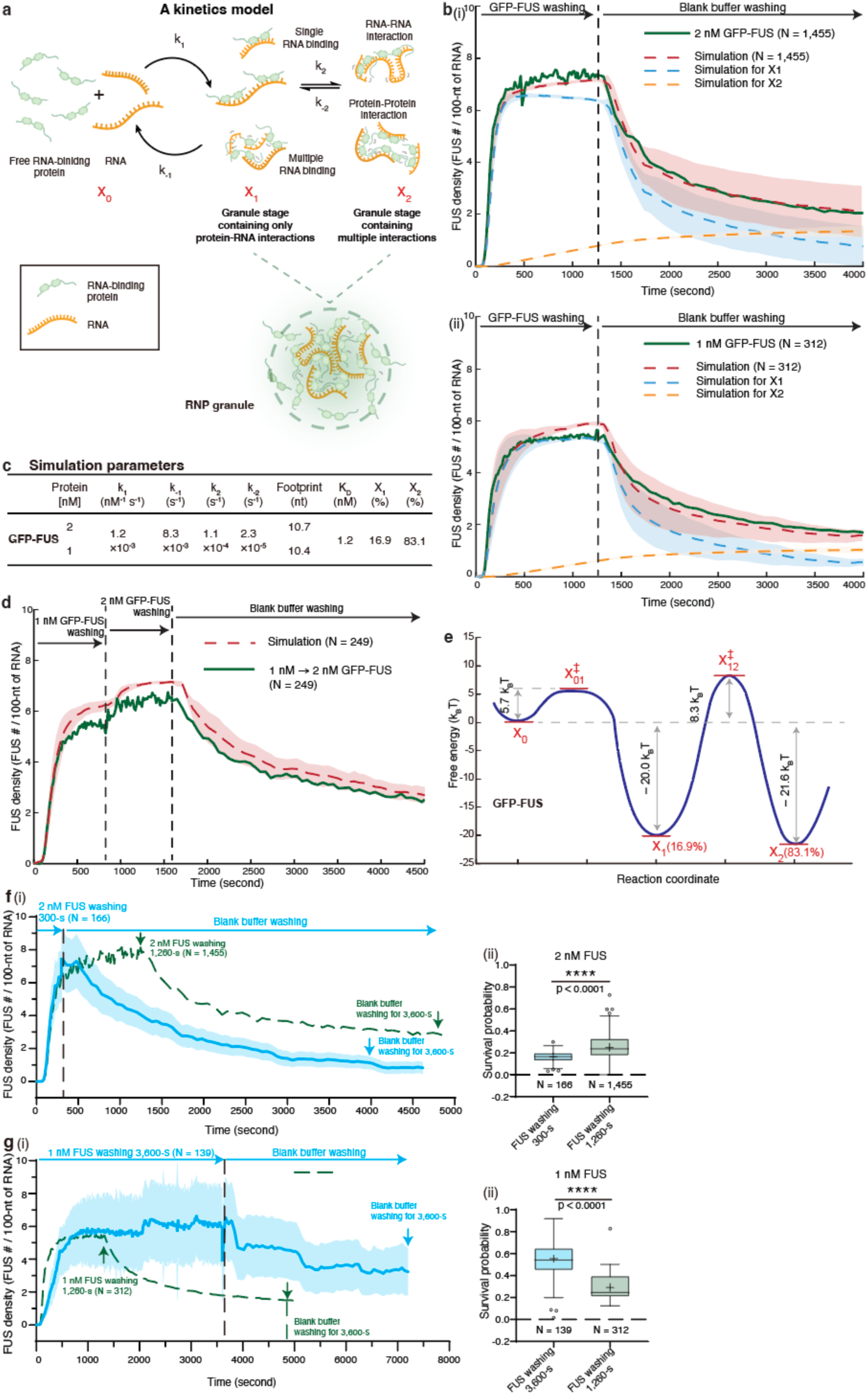
SMART measurements combined with a two-state mathematical model reveal the intrinsically non-equilibrium assembly kinetics of FUS RNP granules. (**a**) Schematic of the two-state mathematical model. (**b**) Simulated SMART measurements during washing following exposure to 2 nM FUS (i) or 1 nM FUS (ii). Red dashed lines indicate simulations to the experimental SMART data; blue and orange dashed lines represent simulated trajectories of state *x*_1_ and state *x*_2_, respectively. (**c**) Table summarizing the simulation parameters. (**d**) Simulation corresponding to the SMART measurement shown in Fig. 1e. All simulations repeat N times, which is the total puncta number of RNP granules examined in the SMART experiments. Simulation results are presented as mean ± s.d. (**e**) Energy landscape of FUS RNP granule assembly inferred from the two-state model. (**f**) (i) SMART measurement of granule assembly dynamics during exposure to 2 nM FUS for 300 seconds (blue), followed by washing with blank working buffer. The green dashed line corresponds to the SMART measurement shown in Fig. 1c(i). N = 166, which was the total puncta number of FUS RNP granules examined over one-time SMART experiment. Data are presented as mean ± s.d. (ii) Boxplot comparing survival probabilities following 300-s and 1,260-s FUS exposure. (**g**) (i) SMART measurement of granule assembly dynamics during exposure to 1 nM FUS for 3,600 seconds (blue), followed by washing with blank working buffer. The green dashed line corresponds to the SMART measurement shown in Fig. 1d. N = 139, which was the total puncta number of FUS RNP granules examined over one-time SMART experiment. Data are presented as mean ± s.d. (ii) Boxplot comparing survival probabilities following 3,600-s and 1,260-s FUS exposure. Survival probability was defined as the ratio of FUS density measured after protein exposure plus 3,600 seconds of blank-buffer washing to the FUS density measured immediately after protein exposure. The total number of granules analyzed in each SMART experiment is denoted by N. The function of “boxplot” in GraphPad Prism software was used to plot the boxplots. For each boxplot, the center line indicates the median; box boundaries represent the 25^th^ and 75^th^ percentiles. Most extreme data points are covered by the whiskers except outliers. Open circles (“o”) is used to represent the outliers. The “+” symbol is used to represent the mean value. Statistical significance was evaluated based on Student’s t-tests (Prism 10 for Windows, Version 10.2.3 (403), April 21, 2024, GraphPad Software, Inc.). Test was chosen as unpaired t test. P value style: GP: 0.1234 (ns), 0.0332 (*), 0.0021 (**), 0.0002 (***), < 0.0001 (****).

Before FUS molecules and RNA substrates assemble into an RNP granule, the system consists solely of freely diffusing components with no detectable inter- or intramolecular interactions; we define this initial condition as state *x*_0_. The power-law disassembly behavior of FUS RNP granules suggests the presence of multiple molecular states supported by distinct interaction types—including RNA-protein, protein-protein, and RNA-RNA contacts. To simplify this complexity on interaction types, we introduce a simplified two-state model. In this framework, the system transitions from the free state *x*_0_ to an intermediate state *x*_1_, in which only protein-RNA interactions are formed. This step is governed by forward and reverse rate constants *k*_1_(nM^-1^×s^-1^) and *k*_-1_ (s^-1^). The intermediate then transitions to a more stabilized state *x*_2_, where additional protein-protein and RNA-RNA interactions emerge. This second step is described by rate constants *k*_2_(s^-1^) and *k*_-2_(s^-1^) (Fig. 2a). Each protein molecule occupies a finite region on the RNA substrate, defined as its *Footprint* (in nt). The *Footprint* parameter may reflect the higher-order architectures adopted by RNA substrates within RNP granules. Together, the four kinetic rate constants and the *Footprint* constitute the five parameters used in the simulations.

We applied this mathematical framework to simulate the assembly and disassembly kinetics under both 2 nM and 1 nM FUS conditions (Fig. 1c-d). The resulting simulations (Fig. 2b(i)-(ii)) matched the experimental data very well. The fitted parameters are summarized in Fig. 2c. Notably, the parameter *Footprint* exhibited comparable values between the 2 nM and 1 nM conditions. Remarkably, the same set of parameters also reproduced the kinetic behavior observed when switching from 1 nM to 2 nM FUS (Fig. 1e), as confirmed by the corresponding simulations (Fig. 2d). Importantly, the sum of simulated rate, *k*_2_ + *k*_-2_, which ∼10^-4^ s^-1^ and determines the timescale over which proteins are retained through cycling between *x*_1_and *x*_2_, is more than two orders of magnitude lower than the corresponding value (∼ 10^#2^ s^-1^) measured in single-molecule assays (*7*). This disparity indicates that multivalent and collective interactions at the mesoscale markedly prolong protein residence times within RNP granules (**Supplementary Methods**).

Previous studies have proposed schematic energy landscapes for RNP granules (*20–22*). Enabled by the SMART assay and our mathematical framework, we can now quantitatively reconstruct the energy landscape of FUS RNP granules (Fig. 2e and **Supplementary Methods**).

### FUS RNP granule assembly kinetics are intrinsically non-equilibrium on experimentally relevent timescales

Analysis of the simulations (Fig. 2c) indicates that thermodynamic equilibrium would require the *x*_2_state to comprise approximately 83.1% of the population. However, under the 2 nM FUS condition, *x*_2_accounts for only ∼11% of the population at the experimental timescale (∼1,260 seconds) (Fig. 2b(i)), demonstrating that FUS RNP granule assembly remains far from equilibrium on experimental timescales. The model further predicts that the characteristic timescale for the 2 nM FUS system (Fig. 2b(i)) to approach equilibrium is ∼1/(*k*_2_ + *k*_-2_) ∼10^4^ seconds, indicating intrinsically slow relaxation dynamics. Consistent with this prediction, simulations based on the fitted parameters show that nearly 40,000 seconds are required for FUS RNP granules to reach their final thermodynamic equilibrium state (Supplementary Fig. 12a).

We hypothesized that varying assembly duration would not substantially alter the saturation density, but that shorter assembly times would yield a less stable, lower-density core architecture. To test this hypothesis, we systematically varied the duration of protein exposure. Reducing the 2 nM FUS assembly time from 1,260-s (Fig. 1c(i)) to 300-s, or extending the 1 nM FUS assembly time from 1,260-s (Fig. 1d) to 3,600-s, resulted in nearly identical saturation densities across all three conditions. In contrast, after an identical 3,600-s blank-buffer wash, the protein density within the stable core architecture of FUS RNP granules was consistently lower following shorter assembly times than after prolonged assembly (Fig. 2f-g). Consistent with these observations, simulations based on the fitted parameters (Supplementary Fig. 12b-c) reproduced this behavior and further revealed that shorter assembly times reduce the population of the *x*_2_state, thereby limiting the formation of the stable core architecture within FUS RNP granules.

Notably, simulations of SMART experiments involving a switch from 1 nM to 2 nM FUS (Fig. 2d), as well as simulations examining variations in assembly duration (Supplementary Fig. 12b-c), quantitatively reproduced the corresponding experimental results without introducing additional parameters. Together, these findings strongly support the validity of the two-state mathematical model.

### SMART reveals general principles of single-component RNP granule biology

Besides FUS, we also quantified the assembly and disassembly kinetics of several additional RBPs, including hnRNPA1a (2 nM and 5 nM in Fig. 3a and Supplementary Fig. 13a-e), G3BP1 (2 nM in Fig. 3b and Supplementary Fig. 14a-d), YTHDF2 (2 nM in Fig. 3c and Supplementary Fig. 15a-d), and TIA1 (2 nM and 5 nM in Fig. 3d and Supplementary Fig. 16). Remarkably, the mathematical model introduced in Fig. 2a simulated the kinetic behaviors of all tested RBPs (Fig. 3e), enabling quantitative reconstruction of the energy landscapes for hnRNPA1a (Supplementary Fig. 13f), G3BP1 (Supplementary Fig. 14e), and YTHDF2 (Supplementary Fig. 15e) RNP granules.

**Fig. 3.**
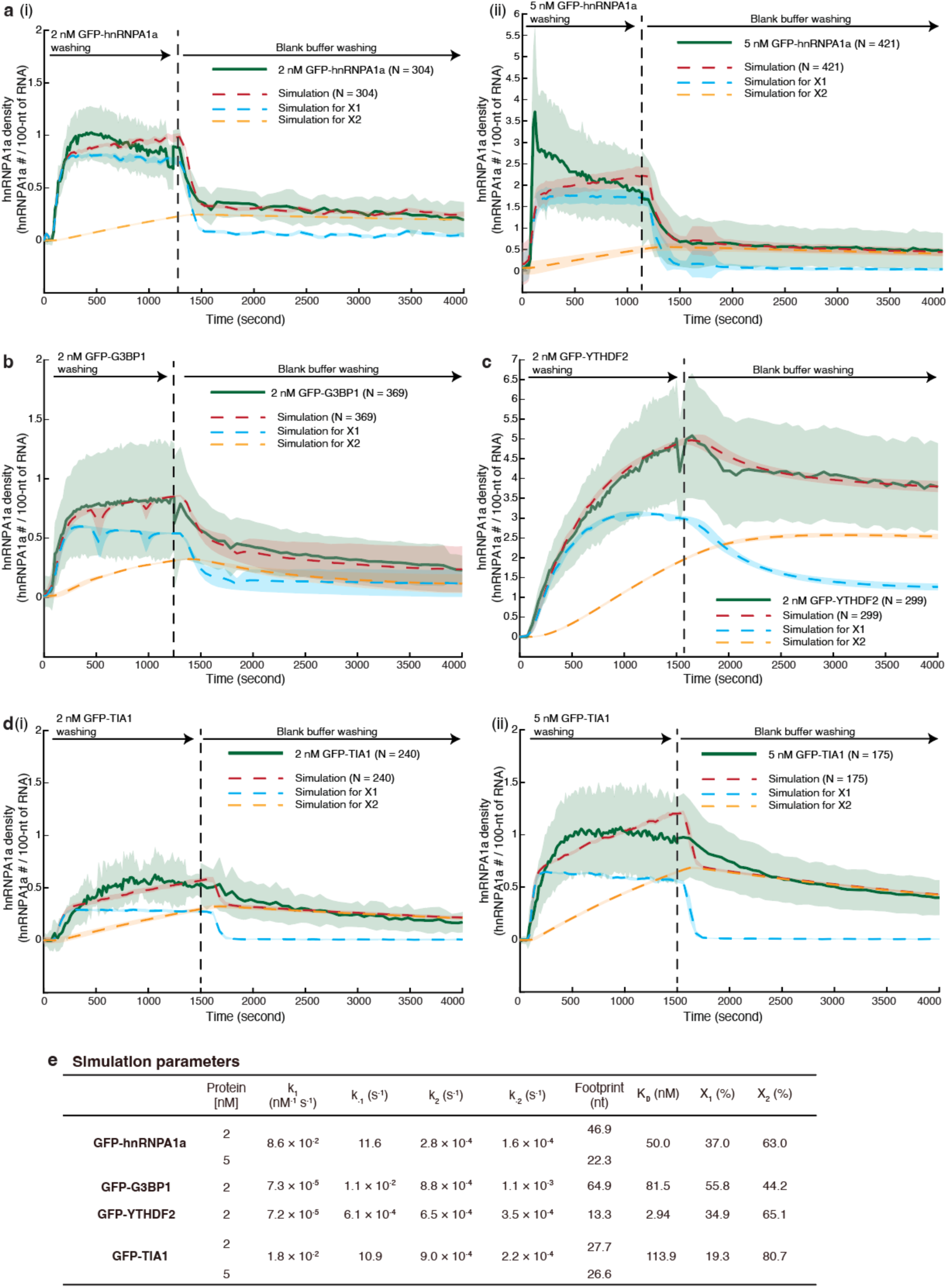
Quantitative SMART measurements of assembly and disassembly kinetics of single-component RNP granules. (**a**) (i) SMART measurement of granule assembly dynamics during exposure to 2 nM GFP-hnRNPA1a for 1,230 seconds, followed by washing with blank working buffer; N = 304, which was the total puncta number of hnRNPA1a RNP granules examined over one-time SMART experiment. (ii) SMART measurement of granule assembly dynamics during exposure to 5 nM GFP-hnRNPA1a for 1,140 seconds, followed by washing with blank working buffer. N = 421, which was the total puncta number of hnRNPA1a RNP granules examined over five times SMART experiments (n = 5). (**b**) SMART measurement of granule assembly dynamics during exposure to 2 nM GFP-G3BP1 for 1,245 seconds, followed by washing with blank working buffer. N = 369, which was the total puncta number of G3BP1 RNP granules examined over three times SMART experiments (n = 3). (**c**) SMART measurement of granule assembly dynamics during exposure to 2 nM GFP-YTHDF2 for 1,530 seconds, followed by washing with blank working buffer. N = 299, which was the total puncta number of YTHDF2 RNP granules examined over three times SMART experiments (n = 3). (**d**) (i) SMART measurement of granule assembly dynamics during exposure to 2 nM GFP-TIA1 (i) or 5 nM GFP-TIA1 (ii) for 1,500 seconds, followed by washing with blank working buffer. N = 240, which was the total puncta number of TIA1 RNP granules (2 nM) examined over two times SMART experiments. N = 175, which was the total puncta number of TIA1 RNP granules (5 nM) examined over one-time SMART experiment. All SMART data are presented as mean ± s.d. Red dashed lines indicate simulations to the experimental SMART data by the two-state mathematical model; blue and orange dashed lines represent simulated trajectories of state *x*_1_and state *x*_2_, respectively. All simulations repeat N times, which is the total puncta number of RNP granules examined in the SMART experiments. Simulation results are presented as mean ± s.d. (**e**) Table summarizing the simulation parameters.

For hnRNPA1a (Supplementary Fig. 13e), raising the concentration from 2 nM to 5 nM increased the saturation level from 0.7 ± 0.2 (mean ± s.d., N = 304 at 1,230 seconds) to 1.8 ± 0.8 (mean ± s.d., N = 421 at 1,140 seconds). Likewise, for TIA1, increasing the concentration from 2 nM to 5 nM elevated the saturation level from 0.5 ± 0.2 (mean ± s.d., N = 240 at 1,500 seconds) to 0.9 ± 0.4 (mean ± s.d., N = 175 at 1,500 seconds) (Supplementary Fig. 16d). These observations further demonstrate that RNP granule kinetics are strongly protein-concentration dependent.

For hnRNPA1a, the *x*_2_state accounted for only 23% of the population at ∼1,230 seconds under 2 nM conditions, whereas thermodynamic equilibrium would require ∼63%. For G3BP1, *x*_2_ comprised 37% at ∼1,245 seconds, compared with an equilibrium requirement of ∼44%. For YTHDF2, *x*_2_ reached only 39% at ∼1,530 seconds, relative to an equilibrium requirement of ∼65%. For TIA1 under 2 nM conditions, *x*_2_reached 52% at ∼1,500 seconds, far below the ∼81% required for equilibrium. Together, these results demonstrate that the assembly of all examined single-component RNP granules proceeds via non-equilibrium pathways on experimental timescales.

### RNA-RBP interactions govern the kinetics of the *x*_1_-state transition

Because the kinetics of FUS RNP granules are governed by nanometer-scale molecular interactions—including RNA-protein, protein-protein, and RNA-RNA interactions—we next examined the contribution of these interactions to granule assembly. FUS comprises two major regions (Supplementary Fig. 5a(i)): a low-complexity domain (LCD) and an RNA-binding domain (RBD). We first purified the FUS LCD (GFP-FUS LCD; Supplementary Fig. 17a(i)-(ii)). Electrophoretic mobility shift assays (EMSAs) revealed minimal RNA-binding activity for the LCD (Supplementary Fig. 17a(iii)). Consistently, SMART measurements showed that GFP-FUS LCD alone failed to assemble into RNP granules (Supplementary Fig. 17b-d), indicating that LCD-LCD interactions are insufficient to drive condensate assembly.

We next purified the FUS RBD (GFP-FUS RBD; Supplementary Fig. 18a(i)-(ii)). EMSA measurements showed that the RBD binds a 30-nt 1× pepper RNA with an affinity of 87.6 ± 8.7 nM (mean ± s.d., n = 3, Supplementary Fig. 18a(iii)-(iv)), comparable to that of full-length FUS (53.0 ± 4.4 nM, mean ± s.d., n = 3; Supplementary Fig. 5b(i)-(ii)). SMART measurements further revealed that the RBD alone compacts RNA and assembles into RNP granules, albeit with slower kinetics than full-length FUS (Fig. 4a and Supplementary Fig. 18b), reaching a protein density of 5.6 ± 1.8 (mean ± s.d., N = 301) at 1,440 seconds. The two-state mathematical model faithfully reproduced these kinetics (Fig. 4a and d), enabling reconstruction of the corresponding free-energy landscape (Supplementary Fig. 18c). Interestingly, simulations indicated that the FUS RBD exclusively populates the *x*_1_ state, with no detectable formation of the *x*_2_state (Fig. 4d). This behavior arises from an ∼8 k_B_T increase in the activation barrier (the free energy between *x*_1_and 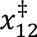 (transition state between *x*_1_ and *x*_2_) in Supplementary Fig. 18c) relative to full-length FUS. Together, these results demonstrate that RNA-FUS interactions are essential for nucleating and stabilizing FUS RNP granules.

**Fig. 4.**
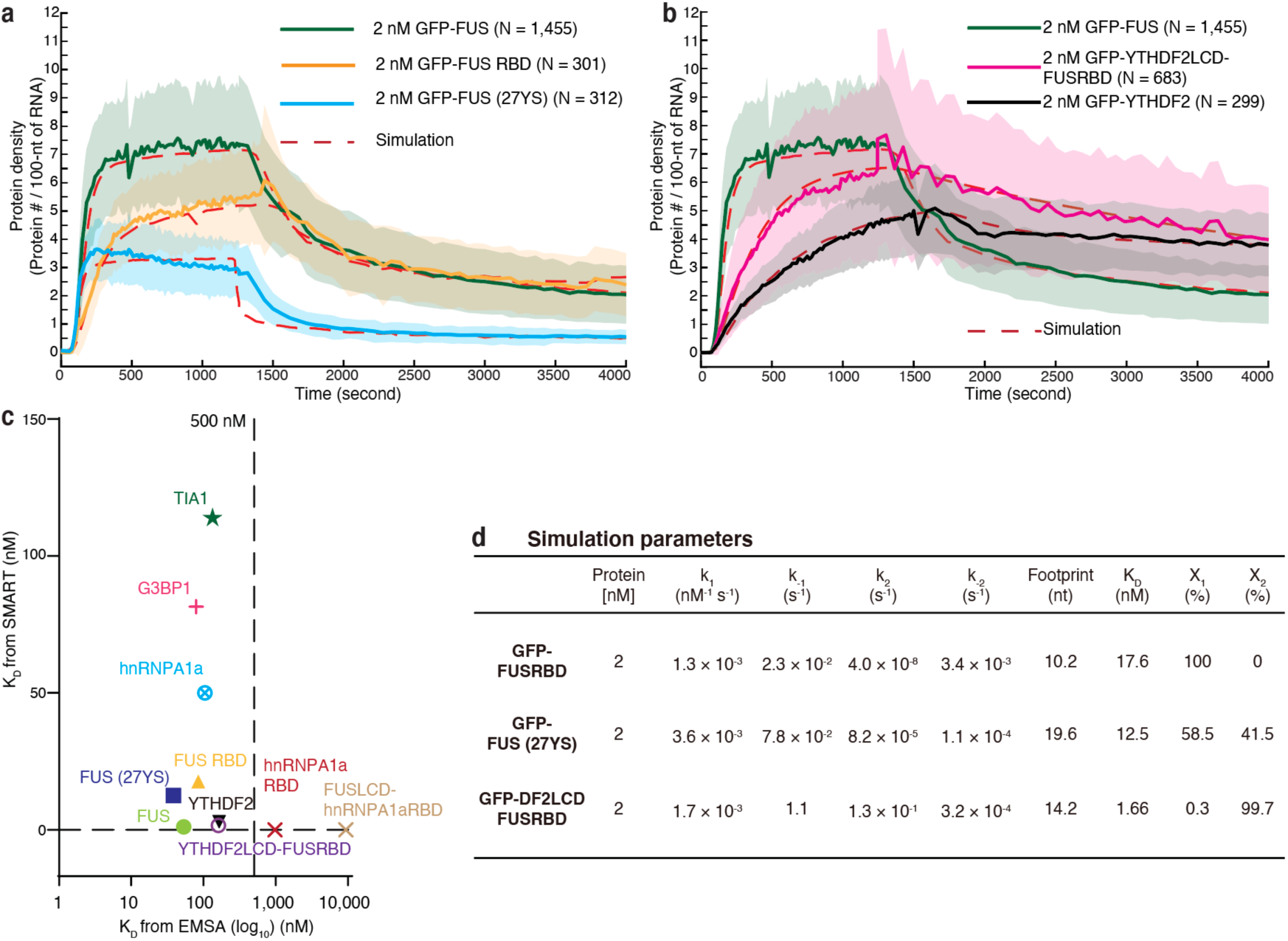
Quantitative SMART measurements of assembly and disassembly kinetics of FUS mutant and truncation RNP granules. (**a**) Green line, SMART measurement of granule assembly dynamics during exposure to 2 nM GFP-FUS for 1,260 seconds, followed by washing with blank working buffer; Orange line, SMART measurement of granule assembly dynamics during exposure to 2 nM GFP-FUS RBD for 1,440 seconds, followed by washing with blank working buffer; N = 301, which was the total puncta number of FUS RBD RNP granules examined over two times SMART experiments. Blue line, SMART measurement of granule assembly dynamics during exposure to 2 nM GFP-FUS (27YS) for 1,260 seconds, followed by washing with blank working buffer. N = 312, which was the total puncta number of FUS (27YS) RNP granules examined over three times SMART experiments (n = 3). (**b**) Green line, SMART measurement of granule assembly dynamics during exposure to 2 nM GFP-FUS for 1,260 seconds, followed by washing with blank working buffer; purple line, SMART measurement of granule assembly dynamics during exposure to 2 nM GFP-YTHDF2LCD-FUSRBD for 1,245 seconds, followed by washing with blank working buffer; N = 683, which was the total puncta number of YTHDF2LCD-FUSRBD) RNP granules examined over four times SMART experiments (n = 4). Black line, SMART measurement of granule assembly dynamics during exposure to 2 nM GFP-YTHDF2 for 1,530 seconds, followed by washing with blank working buffer. All SMART data are presented as mean ± s.d. Red dashed lines in (a)-(b) indicate simulations to the experimental SMART data by the two-state mathematical model. All simulations repeat N times, which is the total puncta number of RNP granules examined in the SMART experiments. (**c**) Dissociation constants derived from EMSAs versus effective dissociation constants extracted from SMART experiments. (**d**) Table summarizing the simulation parameters.

To further probe the role of RNA-RBP interactions, we purified the hnRNPA1a RBD (mCherry-hnRNPA1a RBD; Supplementary Fig. 19a) and a chimeric protein consisting of the FUS LCD fused to the hnRNPA1a RBD (mCherry-FUSLCD-hnRNPA1aRBD; Supplementary Fig. 19c). EMSAs showed that both proteins bind the 30-nt 1× pepper RNA with markedly weaker affinities (>1,000 nM). Consistently, SMART measurements indicated that neither protein formed RNP granules (Supplementary Fig. 19b and d). Interestingly, comparison of dissociation constants derived from EMSAs with effective dissociation constants extracted from SMART experiments (Fig. 4c and **Supplementary Methods**) revealed a threshold affinity (∼500 nM from EMSA), beyond which single-component RNP granule assembly cannot be supported.

Collectively, these results establish RNA-RBP interactions as indispensable determinants of single-component RNP granule nucleation and stability, and identify them as key drivers of the short-dwell-time *x*_1_ state.

### The LCD regulates the kinetics of *x*_2_-state formation

The simulation result that no detectable *x*_2_ -state formation for FUS RBD strongly suggests that the LCD is required to establish and stabilize the *x*_2_ state. To test this hypothesis, we next examined two FUS mutants with perturbed LCD-mediated interactions.

The FUS LCD contains 27 [G/S]Y[G/S] repeats, and previous studies established that tyrosine residues within these motifs are critical for governing FUS’s saturation concentration (*23, 24*). We therefore mutated all 27 tyrosines to serine, generating GFP-FUS (27YS) (Supplementary Fig. 20a(i)-(ii)). EMSA measurements (Supplementary Fig. 20a(iii)-(iv)) showed that its affinity for a 30-nt 1× pepper RNA (36.5 ± 23.4 nM, mean ± s.d., n = 3) is comparable to that of full-length FUS (∼53 nM). This mutant failed to undergo condensation even at 10 μM (Supplementary Fig. 20b), indicating a severe disruption of LCD-LCD interactions. SMART measurements, simulations, and the resulting energy landscape (Fig. 4a, d, and Supplementary Fig. 20c, d, h, i) revealed a markedly reduced protein density (2.9 ± 1.0, mean ± s.d., N = 312 at 1,260-s), compared with ∼7.3 for full-length FUS. Consistent with this reduction, the *x*_2_-state population dropped from 83.1% (full-length FUS) to 41.5%.

In a second perturbation, we replaced the FUS LCD with the LCD from YTHDF2, generating a chimeric protein, YTHDF2LCD-FUSRBD (Supplementary Fig. 20e(i)-(ii)). EMSAs (Supplementary Fig. 20e(iii)-(iv)) revealed that this chimera binds RNA with reduced affinity (197.8 ± 60.8 nM, mean ± s.d., n = 3) compared with full-length FUS (∼53 nM). SMART measurements showed that its assembly kinetics are intermediate between those of full-length FUS and YTHDF2 (Fig. 4b), reaching a protein density of 6.4 ± 3.2 at 1,245 seconds (mean ± s.d., N = 683). Interestingly, the temporal profile of YTHDF2LCD-FUSRBD assembly more closely resembles that of YTHDF2 than that of full-length FUS. Quantitative simulations and reconstruction of the corresponding free-energy landscape (Fig. 4b, d, and Supplementary Fig. 20f-h) revealed that the activation barrier (the free energy between *x*_1_and 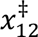 (transition state between *x*_1_ and *x*_2_) in Supplementary Fig. 20g)) for this chimera is reduced by approximately 7 k_B_T relative to full-length FUS, resulting in a markedly increased *x*_2_-state population (99.7%). Together, these results demonstrate that the LCD plays a decisive role in modulating the kinetic barrier and in stabilizing the *x*_2_ state.

### The assembly of two-component RNP granules is pathway dependent

Because endogenous RNP granules comprise multiple RBPs, we next investigated how mixtures of two distinct RBPs assemble and disassemble on shared RNA substrates using the SMART assay. We selected FUS and hnRNPA1a as a representative pair and designed three complementary experimental configurations to systematically probe pathway dependence during two-component RNP granule formation.

In the first configuration, we co-introduced 2 nM GFP-FUS and 5 nM mCherry-hnRNPA1a into the flow cell (Fig. 5a(i) and Supplementary Fig. 21a, d, and e). At 90 seconds, the densities of hnRNPA1a and FUS were 3.2 ± 0.9 (mean ± s.d., N = 79) and 0.2 ± 0.1 (mean ± s.d., N = 79), respectively, indicating that hnRNPA1a initially outcompetes FUS and rapidly occupies the RNA substrate. However, by 210 seconds, the densities shifted to 0.3 ± 0.1 (mean ± s.d., N = 79) for hnRNPA1a and 5.8 ± 1.1 (mean ± s.d., N = 79) for FUS, demonstrating that FUS subsequently displaces hnRNPA1a and dominates the later stages of assembly (Inset of Fig. 5a(i)). Notably, the assembly kinetics of FUS in the mixed-condensate context match those of FUS alone (Fig. 1c(i)).

**Fig. 5.**
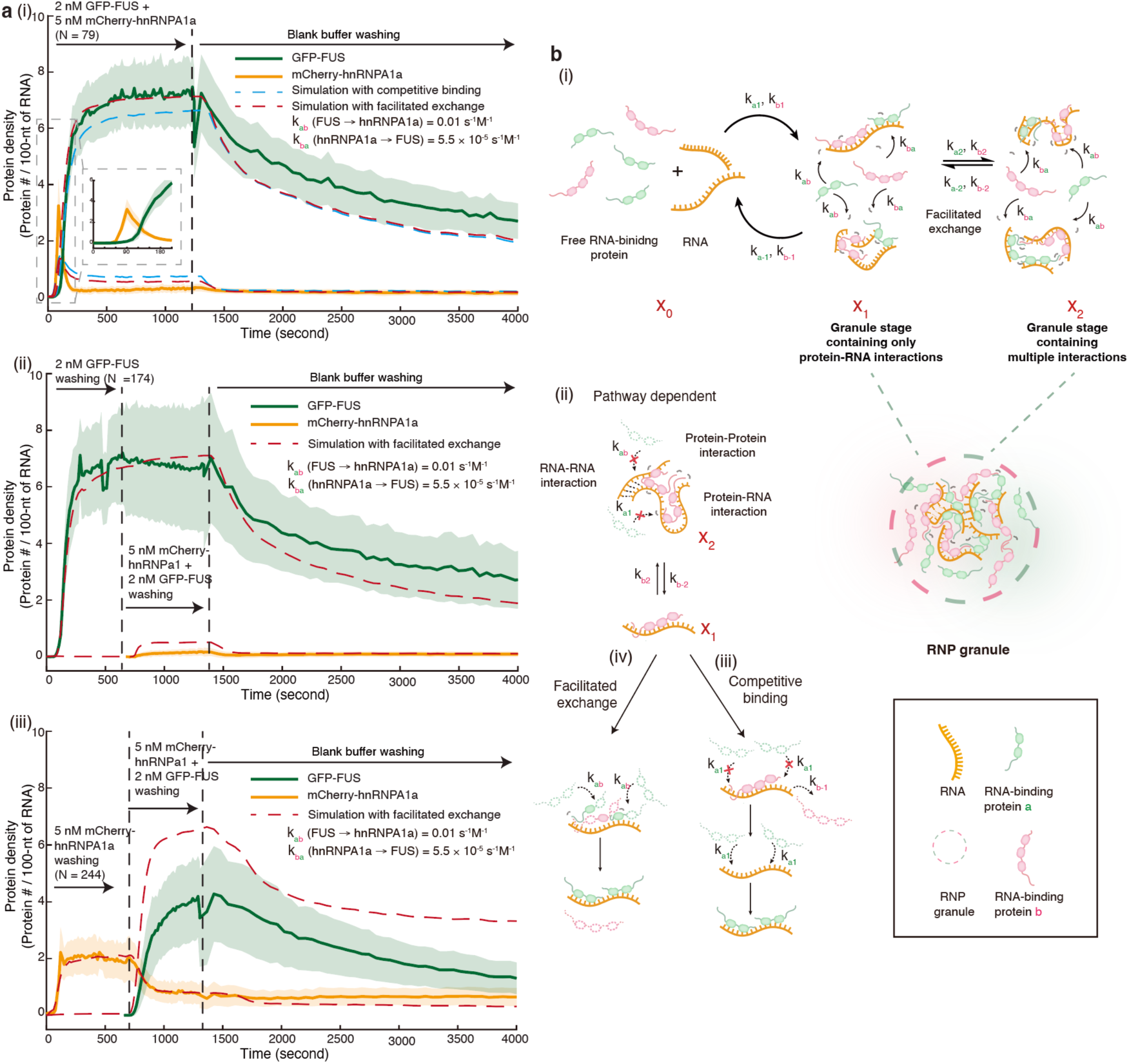
SMART measurements combined with a two-state mathematical model reveal pathway-dependent assembly of two-component RNP granules. (**a**) Three complementary experimental configurations were designed: (i) simultaneous introduction of 2 nM GFP-FUS and 5 nM mCherry-hnRNPA1a into the flow cell; N = 79, which was the total puncta number of FUS-hnRNPA1a RNP granules examined over one-time SMART experiment (1^st^ experimental configuration); (ii) pre-assembly of GFP-FUS for 10 minutes followed by introduction of a mixture of 2 nM GFP-FUS and 5 nM mCherry-hnRNPA1a; N = 174, which was the total puncta number of FUS-hnRNPA1a RNP granules examined over two times SMART experiments (2^nd^ experimental configuration); (iii) pre-assembly of mCherry-hnRNPA1a for 10 minutes followed by introduction of the same protein mixture. N = 244, which was the total puncta number of FUS-hnRNPA1a RNP granules examined over two times SMART experiments (3^rd^ experimental configuration). Green traces indicate GFP-FUS and orange traces indicate mCherry-hnRNPA1a. All SMART data are presented as mean ± s.d. Blue dashed lines represent simulations of the SMART data using the two-state model with a simple competitive-binding hypothesis (b(i)-(iii)). Red dashed lines represent simulations using the two-state model incorporating a facilitated-exchange mechanism (b(i), (ii), and (iv)). All simulations repeat N times, which is the total puncta number of RNP granules examined in the SMART experiments. (**b**) Schematic of the two-state mathematical model with a simple competitive-binding hypothesis (i)-(iii), or a facilitated-exchange mechanism (i), (ii), and (iv).

In the second configuration, we first assembled FUS for 10 minutes and then introduced a mixture of FUS and hnRNPA1a (Fig. 5a(ii) and Supplementary Fig. 21b, d, and e). In this case, hnRNPA1a did not reassemble onto the RNAs upon buffer exchange, and FUS followed its characteristic assembly trajectory, identical to its single-component behavior. This suggests that once FUS gains access to RNAs, hnRNPA1a cannot compete effectively.

In the third configuration, we first assembled hnRNPA1a for 10 minutes and then introduced a mixture of FUS and hnRNPA1a (Fig. 5a(iii) and Supplementary Fig. 21c-e). As expected, hnRNPA1a density decreased after the addition of FUS, confirming that FUS can compete hnRNPA1a off the RNAs. However, FUS reached a density of only 4.0 ± 1.4 (mean ± s.d., N = 244)—substantially lower than ∼7.3 observed for 2 nM FUS alone (Fig. 1c(i)). Thus, preassembled hnRNPA1a impedes the complete assembly of FUS. Moreover, nearly all hnRNPA1a molecules retained within the granule contribute to the formation of the final stable core structure of hnRNPA1a RNP granules.

Together, these results demonstrate that the assembly of FUS-hnRNPA1a RNP granules is strongly pathway dependent. We propose that proteins that assemble first have sufficient time to establish stable interactions (*x*_2_-state). Proteins in this state require a longer reaction time to expose their RNA-binding sites (Fig. 5b(ii)). Similar behavior was observed for FUS-G3BP1 (Supplementary Fig. 22) mixtures, indicating that pathway dependence is a general principle governing the assembly of multi-RBP RNP granules.

### Our mathematical model incorporating a facilitated-exchange term better simulates the kinetics of two-component RNP granules

We next examined whether our mathematical framework could capture the assembly dynamics of FUS-hnRNPA1a RNP granules. As a starting point, we directly combined the independently derived kinetic parameters for FUS and hnRNPA1a RNP granules (Fig. 2c and 3e) and proposed a simple competitive-binding hypothesis (Fig. 5b(i)-(iii)). In this scenario, if a given RNA segment is occupied by one RBP, a second RBP can bind only after the first dissociates. Using this scheme, we simulated the SMART experiment in which 2 nM GFP-FUS and 5 nM mCherry-hnRNPA1a were co-introduced into the flow cell (Fig. 5a(i)). However, the resulting simulations (blue dashed lines) failed to reproduce the experimental kinetics for either protein.

To address this discrepancy, we incorporated a facilitated-exchange mechanism, inspired by work from Marko and co-workers showing that freely diffusing DNA-binding proteins can accelerate the dissociation of DNA-bound counterparts (*25*). We hypothesized (Fig. 5b(iv)) that FUS can facilitate the dissociation of hnRNPA1a (rate constant **k*_FUS→,hnRNPA1a_*, s^-1·^M^-1^), and conversely, that hnRNPA1a facilitates the dissociation of FUS (**k*_hnRNPA1a_*_→*FUS*_, s^-1·^M^-1^). We hypothesize that this displacement effect is applicable to both states. Introducing these two additional parameters improved the agreement between simulations and experiments (red dashed lines in Fig. 5a(i)), providing a reasonable basis for the facilitated-exchange hypothesis.

Interestingly, the simulated facilitated-exchange parameters from the first experimental configuration (Fig. 5a(i)) also reproduced the kinetics in the second configuration (Fig. 5a(ii)), but failed to account for the third configuration (Fig. 5a(iii)). This divergence may suggest two possible mechanisms for pathway dependence. First, the displacement rate may differ for proteins in the two states, or that proteins in the *x*_2_ state cannot be displaced at all (Fig. 5b(ii)). Second, the pre-assembled hnRNPA1a may produce different underlying RNA-RNA architectures within two-component RNP granules, resulting in pathway-specific facilitated-exchange dynamics.

## Discussion

In this study, we developed a new single-molecule technique—SMART, which can be used to directly monitor the cross-scale kinetics of RNP granules from nanometer-scale interactions, such as RNA-protein, protein-protein, and RNA-RNA interactions, to the assembly of mesoscale RNP granules. We also established a two-state mathematical model to successfully simulate the assembly and disassembly kinetics of RNP granules. During the process of assembly, the initial state *x*_0_containing no any biomolecular interactions firstly transitions to state *x*_1_, which is dominated by RNA-RBP interactions; *x*_1_then secondly transitions to state *x*_2_, which is regulated by the LCD of RBP. Besides the RNA-RBP interactions, additional protein-protein and RNA-RNA interactions emerge in state *x*_2_.

SMART, together with the two-state mathematical model, was first applied to investigate single-component RNP granules formed by individual RBPs, including FUS, hnRNPA1a, G3BP1, YTHDF2, and TIA1, enabling quantitative reconstruction of the energy landscape for each system. Although different RBPs exhibited distinct assembly and disassembly kinetics, the assembly of all single-component RNP granules was inherently non-equilibrium. Extending this framework to two-component RNP granules revealed that their assembly kinetics are strongly pathway dependent. Consistent with our findings, starvation- and heat-induced stress granules in budding yeast display distinct dissolution kinetics of the RNA-binding protein Pub1 (*26*), suggesting that stress granule assembly in living cells is also pathway dependent.

Our findings identify several general principles of RNP granule biology that motivate future investigation. First, the intracellular concentrations of most RBPs in vivo are substantially lower than their corresponding saturation concentrations measured in vitro (*24, 27*). For instance, the cellular concentrations of FUS and hnRNPA1a are approximately 2.3 μM and 12.7 μM, respectively, whereas the saturation concentration of FUS and many other RBPs in vitro ranges from ∼5 to 100 μM. Recently, Grill and colleagues applied adsorption and prewetting theory—well-established concepts from soft-matter physics—to demonstrate that the transcription factor Klf4 can form biomolecular condensates on DNA surfaces at concentrations below its bulk saturation threshold (*28*). These findings raise the intriguing possibility that RBPs may similarly undergo adsorption and prewetting on RNA substrates, thereby enabling RNP granule formation at sub-saturation concentrations. Exploring this mechanism represents a promising avenue for future research.

Second, our SMART experiments in which 1 nM FUS was exchanged with 2 nM FUS (Fig. 1e) suggest that RNA within RNP granules remains partially accessible. An analogous concept is well established for double-stranded DNA condensates in the context of chromatin accessibility, wherein not all genomic regions are continuously occupied by DNA-binding proteins (*29*). Recently, we further demonstrated that replication protein A (RPA), the primary single-stranded DNA-binding protein in eukaryotes, cannot fully occupy extended ssDNA substrates over long timescales, revealing a form of ssDNA accessibility (*30*). Taken together, these observations suggest that “accessibility” may represent a general principle of biomolecular condensates formed on long nucleic acid substrates. Elucidating the molecular mechanism of nucleic acid accessibility within RNP granules

Third, for a subset of RBPs, our SMART experiments suggest that multivalent interactions within RNP granules generate a pronounced separation of timescales corresponding to the *x*_1_and *x*_2_states. The *x*_1_state, characterized by a short timescale (∼10^2^ seconds), reflects rapid protein binding and dissociation, whereas the *x*_2_ state, operating on a much longer timescale (∼10^4^ seconds), captures stabilized multivalent interactions mediated by protein-protein and RNA-RNA contacts. This separation of timescales may constitute a general principle governing RNP granule assembly. Protein-specific differences in the transition rates between these states define a regulatory window for downstream compositional control. At the macroscopic level, this behavior is manifested as markedly distinct diffusion properties of individual protein components within stress granules and processing bodies (*31–33*). More broadly, timescale separation in mesoscopic condensates may have important implications for disease pathogenesis and warrants further investigation.

Fourth, SMART experiments in which preassembled hnRNPA1a granules (5 nM) were challenged with a mixture of 5 nM hnRNPA1a and 2 nM FUS (Fig. 5a(iii)) revealed that pre-existing hnRNPA1a assemblies impede the full incorporation of FUS. One plausible explanation is the formation of pathway-dependent RNA-RNA interaction networks. Specifically, initial hnRNPA1a RNP granules may establish distinct RNA-RNA interaction architectures. Subsequent assembly of hnRNPA1a-FUS granules then proceeds on this preconfigured RNA network, resulting in a final architecture characterized by a lower FUS density (∼4) than that achieved by direct assembly with 2 nM FUS alone (∼7.2; Fig. 1c(i)). Consistent with this interpretation, Parker and colleagues recently demonstrated that G3BP1 promotes the formation of intermolecular RNA-RNA interaction networks (*17, 34*). More broadly, RNA is increasingly recognized as a central structural and functional component of many RNP granules (*1, 35*). Indeed, RNA molecules, rather than proteins, catalyze key steps in pre-mRNA splicing, and ribosomal RNAs, rather than proteins, take charge of the ribosome’s overall structure. Together, these observations motivate future investigations into whether specific intermolecular RNA-RNA interaction networks within RNP granules encode distinct biological functions.

Deciphering the biological functions of complex systems requires quantitative measurement of assembly and disassembly kinetics across multiple length scales. Although this remains a formidable challenge, recent studies have begun to address it. For example, Brangwynne and colleagues developed a suite of approaches to elucidate pre-rRNA processing by bridging nanometer-scale biomolecular interactions within the nucleolus to its mesoscale organization (*3*). Here, we anticipate that the newly developed SMART platform, together with the two-state mathematical model, will directly bridge nanometer-scale molecular interactions and higher-order mesoscale assembly of RNP granules. By enabling quantitative, cross-scale kinetic measurements, this framework provides a powerful approach for decoding the biological functions of RNP granules and may ultimately inform the development of new therapeutic strategies for human disease.

## Author Contributions

Y.L. prepared biological samples, designed and conducted biochemical and single-molecule experiments, performed data analysis, and wrote the manuscript. X.L. conducted theoretical simulation. M.X. assisted Y.L. Z.Q. supervised the project, experimental designs, and data analysis, and wrote the manuscript with input from all authors.

## Acknowledgments

We thank the Peking Nanofab for process support. We thank the contributions of the Engineering Research Center for Semiconductor Integrated Technology, Institute of Semiconductors, Chinese Academy of Sciences. We thank Pilong Li laboratory (Tsinghua University, China) for their generous support of plasmid of YTHDF2. We thank Wei Wang laboratory (Peking University, China) for their generous support of plasmid of TIA1. We thank Dr. Luhua Lai and Jun Liu (Peking University), Dr. Chunlai Chen and Pilong Li (Tsinghua University), and Dr. Cheng Li and other members of the Z.Q. laboratory for comments on the manuscript.

## Methods

### Construction of bacterial expression plasmids

FUS, hnRNPA1a, G3BP1 genes came from human cDNAs. Plasmid of YTHDF2 was a gift from Pilong Li Lab (Tsinghua University, China). Plasmid of TIA1 was a gift from Wei Wang Lab (Peking University, China). The gene of MS2 coat protein was obtained from Addgene (Plasmid nos. 103831). The vector for bacterial protein expression was pRSF-Duet (Novagen). For all proteins, we modify this vector to prepare a 7× His-tagged vector (His vector), a 7× His-GFP-tagged vector (GFP vector), a 7× His-mCherry-tagged vector (mCherry vector). Only protein G3BP1 contains a MBP tag followed with a HRV 3C cleavage site between His tag and GFP/mCherry. All tags and proteins were fused with a 4xGGS linker. The plasmid of protein FUS (27YS) mutant, FUS RBD, hnRNPA1a RBD and two fusion proteins YTHDF2 LCD-FUS RBD, FUS LCD-hnRNPA1a RBD were modified based on the vectors mentioned above. The GFP protein in all constructs is the superfolder GFP (*36*) with a single mutation (A206K) to avoid the weak dimerization.

### Protein purification

GFP and mCherry versions of the same protein is purified with identical protocol. Protein FUS, FUS (27YS), FUS LCD, FUS RBD and fusion protein YTHDF2 LCD-FUS RBD, FUS LCD-hnRNPA1a RBD are purified based on the previous protocol (*7*). Plasmids are transformed into *E. coli* strain BL21 (DE3). Monoclone is inoculated into 5ml LB and incubated overnight. Transfer the 5 mL culture into 1L LB and then shake until the OD reach 0.6-0.8 under 37 °C, 220 rpm. Cultures are shifted into 16 °C, 160 rpm, after adding 300 mM IPTG and incubated overnight. Cells are centrifuged at 3,000 g and re-suspended with lysis buffer (50 mM Tris-HCl (pH 7.5), 1 M KCl, 1 M Urea, 0.1 mg/mL RNase A, 5% (v/v) glycerol, 5 mM β-Mercaptoethanol (βME), 1 mM PMSF (phenylmethyl-sulfonyl fluoride), and 10 mM imidazole-HCl (pH 7.5)) and then sonicated for 10 minutes with 8 seconds sonication plus 16 seconds pulses at 40% amplitude. The supernatant is collected after centrifugation at 18,000 g for 15 minutes. Load the supernatant throw a gravity-flow column filled with 5-10 mL Ni-NTA resin (Thermo Scientific™) and wash the column with 100-150 mL (20-30 column volume) washing buffer (50 mM Tris-HCl (pH 7.5), 1 M KCl, 1 M Urea, 5% (v/v) glycerol, 5 mM βME, 40 mM imidazole-HCl (pH 7.5)). Protein is eluted with elution buffer (50 mM Tris-HCl (pH 7.5), 1 M KCl, 1 M Urea, 5% (v/v) glycerol, 5 mM βME, 300 mM imidazole-HCl pH 7.5) and concentrated using Amicon Ultra-15 Centrifugal Filter 50 kDa or 30 kDa MWCO. The concentrated samples are further filtered (0.22 μm), purified with a Superdex 200 column (GE Healthcare, USA) pre-equilibriumed with storage buffer (50 mM Tris-HCl (pH 7.5), 1 M KCl, 1 M Urea, 5% (v/v) glycerol, 1 mM DTT). Fractions are analyzed by 10% SDS-PAGE and concentrated as above. The aliquots are flash frozen in liquid nitrogen and stored at -80 °C.

Purification of hnRNPA1a, TIA1, YTHDF2 are quite same as above with a minor difference on the buffer. For hnRNPA1a and TIA1, all purification buffer contains no urea and the final storage buffer contains only 500 mM KCl. For protein YTHDF2, the storage buffer contains 40 mM HEPES (pH 7.5), 250 mM KCl, 5 mM DTT and 5% Glycerol.

Purification of G3BP1 is based on the previous protocols (*17, 37*). The construct contains 7× His tag followed with MBP-GFP/mCherry tag at the N-terminal of G3BP1 with a HRV 3C cleavage site between MBP and GFP/mCherry tag. The plasmid is transformed into Rosetta 2 (DE3) strain and cultured, purified the same way as above using Ni-NTA column. The lysis buffer contains 50 mM HEPES-NaCl (pH 7.5), 1 M NaCl, 5 mM βME, 0.1 mg/mL RNase A, 1× Protease inhibitor cocktail (Beyotime, P1005), 1 mM PMSF. The wash buffer contains 50 mM HEPES-NaCl (pH 7.5), 100 mM NaCl, 5 mM βME, and 35 mM imidazole-HCl (pH 7.5). The elution buffer contains 50 mM HEPES-NaCl (pH 7.5), 100 mM NaCl, 5 mM βME, 250 mM imidazole-HCl (pH 7.5). Concentrate the elution sample and exchange the buffer into Buffer A (50 mM HEPES-NaCl (pH 7.5), 100 mM NaCl, and 1 mM DTT) using Amicon Ultra-15 Centrifugal Filter 50 kDa MWCO and gently pipette the solution every 10 minutes to disrupt aggregation as mentioned (*17*). The sample is incubated with RNase A and HRV 3C protease (Sangon Biotech) at 4 °C overnight to remove MBP tag. The proteins are further purified with a HiTrap SP column re-equilibriumed with Buffer A. A gradient elution is performed with Buffer B (50 mM HEPES-NaCl (pH 7.5), 1 M NaCl, 1 mM DTT) ranging from 0%-100% within 10 column volumes. The right samples are concentrated based on SDS-PAGE. The proteins are then purified with Superdex 200 column (GE Healthcare, USA) with storage buffer (50 mM HEPES-NaCl (pH 7.5), 400 mM NaCl, and 1 mM DTT). The final aliquots are flash frozen in liquid nitrogen and stored at -80 °C.

The concentration is calculated with Nanodrop spectrophotometer using Lambert-Beer’s law, for GFP (absorbance at 488 nm, Ec: 83,300 M^-1^cm^-1^) and mCherry (absorbance at 587 nm, Ec: 72,000 M^-1^cm^-1^).

### Electrophoretic mobility shift assay (EMSA)

#### Single-stranded RNA (ssRNA) binding

EMSA experiments are conducted to test the RNA binding affinity of different RNA binding proteins. All experiments are prepared in the same working buffer (40mM Tris-HCl (pH 7.5), 150 mM KCl, 1 mM DTT, 2 mM MgCl_2_, 0.5 mg/mL BSA, and 2 unit/μL RNasin® Plus RNase Inhibitor) by mixing the protein with 10 nM Cy5 labeled RNA substrates at different molar ratios. The total reaction volume is 10 μL. The mixture is incubated at room temperature for 30 mins and then loaded on a 6% native PAGE (polyacrylamide gel electrophoresis) gel running with 0.5 x TBE buffer, 150 V, 30 minutes. The gel is imaged Amersham Typhoon RGB. The binding affinity was fitting with a hill function.

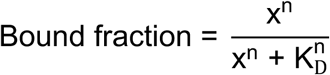

#### ssRNA substrate sequences

5’ – Cy5 – UUU CCC AAU CGU GGC GUG UCG GCC UCU UUU – 3’ (30-nt 1× pepper)

### In vitro Droplet assay

All proteins are centrifuged at 10,000 g for 10 minutes at 4 °C before usage. The proteins are then diluted into various concentrations with BSA working buffer. The total volume is 10 μL. The final system contains 40mM Tris-HCl (pH 7.5), 150 mM KCl, 1 mM DTT, 2 mM MgCl_2_, and 0.5 mg/mL BSA. The sample is transferred into 384-well plates (Cellvis) and incubated at room temperature for 30 mins. The sample is imaged with confocal microscopy (LeicaTCS SP8).

### Bulk biochemical assay for *in vitro* transcription

The reaction is conducted in a 100 μL solution with ∼1 μg dsDNA template, 10 μL 10× reaction buffer (NEB), 1 μL T7 RNA polymerase (NEB), 1 mM NTP mix, 2 unit/μL RNasin® Plus RNase Inhibitor. The mixture is incubated at 37 °C for 4-5 hours. Add 1 μL RNase free DNase I to digest dsDNA template for 1 hour. Add 0.5 μL Protease K (NEB) and incubate for another 30 minutes. Either use RNA purification kit (TIANGEN) to extract RNA substrates or direct load the mixture on to a 1% agarose gel.

### Labeling efficiency of 647-X-UTP

The labeling efficiency is evaluated by conducting a series bulk in vitro transcription reaction as above and varying the incorporation concentration of 647-X-UTP. The dsDNA template is 221 bps which can generate a length of 197 nts RNA product containing 54 U in total. The final concentration of RNA substrate purified with RNA purification kit is calculated based on the absorbance at 260 nm with Ec of 1951200 M^-1^cm^-1^ ([RNA], μM). The concentration of 647-X-UTP is based on the absorbance at 650 nm with Ec of 250,000 M^-1^cm^-1^ ([647-X-UTP], μM). The concentration of NTP is defined by absorbance at 260 nm (A_260_).

The initial incorporation ratio of transcription mix “m_0_” is defined by [647-X-UTP]_0_ (μM) and 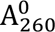 for NTP. The final labeling efficiency “η_0_” is defined by incorporated [647-X-UTP]_final_ (μM) and [RNA] (μM) after RNA purification with kit.

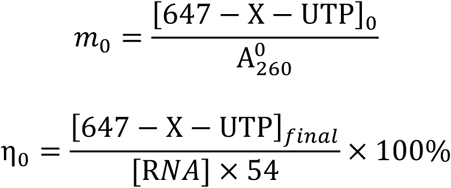

For each *in vitro* transcription reaction, we can plot the distribution of “m_0_-η_0_” and perform a linear fitting. The slope is the labeling coefficient “k”. The labeling efficiency for each SMART experiment “η” is adjusted as the formular below.

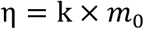

#### RNA sequence

5’-GGUGAUAAGUGGAAUGCCAUGGUUUUAGAGCUAGAAAUAGCAAGUUAAAAUAAGGCUAGUCCGUUAUCAACUUGAAAAAGUGGCACCGAGUCGGUGCUUUUUUUGAAUUCACUGGCCGUCGUUUUACAACGUCGUGACUGGGAAAACCCUGGCGUUACCCAACUUAAUCGCCUUGCAGCACAUCCCCCUUUCGCC AG-3’

#### dsDNA template

5’-TATGACTAATACGACTCACTATAGGGTGATAAGTGGAATGCCATGGTTTTAGAGCTAGAAATAGCAAGTTAAAATAAGGCTAGTCCGTTATCAACTTGAAAAAGTGGCACCGAGTCGGTGCTTTTTTTGAATTCACTGGCCGTCGTTTTACAACGTCGTGACTGGGAAAACCCTGGCGTTACCCAACTTAATCGCCTTGCAGC ACATCCCCCTTTCGCCAG-3’

### Unpaired t test

Statistical significance was evaluated based on Student’s t-tests (Prism 10 for Windows, Version 10.2.3 (403), April 21, 2024, GraphPad Software, Inc.). Test was chosen as unpaired t test. P value style: GP: 0.1234 (ns), 0.0332 (*), 0.0021 (**), 0.0002 (***), < 0.0001 (****).

### Box-plot

The function of “boxplot” in GraphPad Prism software was used to plot the boxplots. For each boxplot, the center line indicates the median; box boundaries represent the 25^th^ and 75^th^ percentiles. Most extreme data points are covered by the whiskers except outliers. Open circles (“o”) is used to represent the outliers. The “+” symbol is used to represent the mean value.

